# IRES-mediated ribosome repositioning directs translation of a +1 overlapping ORF that enhances viral infection

**DOI:** 10.1101/303388

**Authors:** Craig H Kerr, Qing S Wang, Kyung-Mee Moon, Kathleen Keatings, Douglas W Allan, Leonard J Foster, Eric Jan

**Author notes:** Contributed equally (Craig H Kerr, Qing S Wang).

## Abstract

RNA structures can interact with the ribosome to alter translational reading frame maintenance and promote recoding that result in alternative protein products. Here, we show that the internal ribosome entry site (IRES) from the dicistrovirus Cricket paralysis virus drives translation of the 0-frame viral polyprotein and an overlapping +1 open reading frame, called ORFx, via a novel mechanism whereby a subset of ribosomes recruited to the IRES bypasses downstream to resume translation at the +1-frame 13th non-AUG codon. A mutant of CrPV containing a stop codon in the +1 frame ORFx sequence, yet synonymous in the 0-frame, is attenuated compared to wild-type virus in a *Drosophila* infection model, indicating the importance of +1 ORFx expression in promoting viral pathogenesis. This work demonstrates a novel programmed IRES-mediated recoding strategy to increase viral coding capacity and impact virus infection, highlighting the diversity of RNA-driven translation initiation mechanisms in eukaryotes.

**Significance Statement:** Viruses use alternate mechanisms to increase the coding capacity of their viral genomes. Here, we provide biochemical evidence that ribosomes recruited to the dicistrovirus cricket paralysis virus IRES undergo a bypass event to direct translation of a downstream +1 frame overlapping open reading frame, called ORFx. Mutations that block ORFx expression inhibit +1 frame translation and infection in fruit flies. These findings highlight the diversity of RNA-driven translation initiation mechanisms in eukaryotes.

## Introduction

The ribosome mediates translation involving decoding the open reading frame codon by codon through delivery of the correct aminoacyl-tRNAs to the ribosomal A site. This fundamental process occurs with high fidelity for proper gene expression in all species. However, mechanisms exist that can alter the translational reading frame, thus producing alternative protein products from a single RNA (1). In general, these mechanisms termed recoding, involve a specific RNA structure or element that interacts with the ribosome to cause the translating ribosome to shift reading frame by -1/+1/+2 allowing it to read through stop codons or bypass sequences and restart translation downstream (2–4). Study of these mechanisms has been enlightening; revealing key ribosome:RNA interactions that alter fundamental processes in the mechanics of ribosome decoding and reading frame maintenance. Importantly, recoding mechanisms are now appreciated as important regulatory processes that can impact the fate of protein expression in cells and viral infections (1, 5). Unlike these recoding mechanisms that involve an actively translating ribosome, the intergenic internal ribosome entry site (IRES) within a subset of dicistroviruses has the unusual property to directly recruit the ribosome and initiate translation from overlapping 0 and +1-frame codons to produce two distinct proteins (6). Here, we report a novel recoding mechanism and translational initiation pathway whereby a related dicistrovirus IRES directs the ribosome to initiate translation downstream.

Most eukaryotic mRNAs utilize a cap-dependent scanning mechanism involving >12 translation initiation factors to recruit the ribosome and initiate translation from an AUG start codon (7). Alternatively, an IRES, in general, is a structured RNA element that facilitates 5’ end-independent translation using subsets of translation initiation factors, thus providing an advantage during viral infection or under times of stress when cap-dependent translation is compromised (3–5).

Of the different classes of viral IRESs based on factor requirement and mechanism, the intergenic IRES of the *dicistroviridae* family stands out as the most streamlined using a unique mechanism where it directly binds 40S and 80S ribosomes without the need for canonical initiation factors or initiator Met-tRNA_i_ and initiates translation from a non-AUG codon (8–12). The dicistrovirus IRES is composed of three pseudoknots (PKI, II, and III) that separate into distinct domains; PKII and PKIII fold independently to create the ribosome-binding domain while PKI mediates positioning of the ribosome and establishes the translational reading frame (8, 10, 13). Structural studies have indicated that PKII and PKIII form a compact core structure, and the PKI region adopts a conformation that mimics an anti-codon:codon interaction that binds the conserved core of the ribosome in the A site (14–17). From here, in an elongation factor 2 dependent manner, the IGR IRES pseudo-translocates to the ribosomal P site, followed by aminoacyl-tRNA delivery to the A site and a second round of eEF2-dependent pseudo-translocation of the IGR IRES to the E site of the ribosome (17–19). Altogether, the IGR IRES acts as a complete RNA machine that supersedes initiation factors and commandeers the ribosome, a strategy that is essential for viral protein synthesis in dicistrovirus-infected cells (20).

In general, the dicistrovirus IGR IRESs are conserved at the structural, but not sequence level and are classified into two sub-groups (termed Type I and II) based on the presence of distinct structural elements; the main distinction comes from a larger L1.1 loop and an additional stem-loop (SLIII) in Type II IRESs (21, 22). SLIII allows for the PKI domain of Type II IRESs to mimic the global shape of a tRNA in addition to assisting in reading frame selection and the larger L1.1 region functions to mediate 60S recruitment (15, 18, 23). The domains of Type I and II IGR IRESs function similarly to directly recruit 80S ribosomes and initiate translation (24–27). Recent high resolution cryo-EM structures of the IGR IRES bound to the 80S ribosome have demonstrated that the IRES initially binds in the A site (17, 19, 28). Movement of the IRES involves an eEF2-dependent pseudo-translocation event where the ribosome rotates up to 10° allowing PKI to move into the P site in an inchworm-like manner (28). This allows for the non-AUG initiation codon of the IRES to be presented in the A site for the incoming amino-acyl tRNA. The first pseudo-translocation event and delivery of the first amino-acyl tRNA are the rate-limiting steps of initiation on the IGR IRES (29).

Biochemical, phylogenetic, and bioinformatics analyses have demonstrated that a subset of Type II IGR IRESs can direct translation of a hidden +1 open reading frame (ORF), termed ORFx, within ORF2 of the viral genome (6, 30). The functional role of ORFx during viral infection remains elusive. Extensive mutagenesis of the PKI domain of the *Israeli acute paralysis virus* (IAPV) IGR IRES has revealed that 0 and +1 frame translation can be uncoupled, suggesting that the IGR IRES may adopt specific conformations that govern the translational reading frame (18, 31). Generally, Type I and II IGR IRESs are thought to operate similarly in mechanism. Specific domains between the two types are functionally interchangeable (24). In the present study, we investigate the capacity of other IGR IRESs from dicistroviruses to facilitate +1-frame translation. We show that the IGR IRES from Cricket paralysis virus (CrPV) can synthesize an ORFx protein using an unexpected mechanism that involves IRES-mediated ribosome bypassing. Furthermore, we provide insight into the role of ORFx during CrPV infection and show that mutants deficient in ORFx have impaired virulence in adult flies, thus uncovering a novel viral recoding strategy that is essential for viral infection.

## Results

### IGR IRES-dependent +1-frame translation is not conserved throughout *Dicistroviridae*

We previously showed that a subset of dicistrovirus IGR IRESs can direct translation in the 0- and +1-frames and that a base pair adjacent to the PKI domain is important for initiation in the +1-frame (6, 31). The IGR IRESs are classified into two types: Type I, and II, with the main difference being an extra SLIII within the PKI domain of Type II IRES. Since the honeybee and fire ant viruses harbour Type II IGR IRESs that can support +1-frame ORFx translation, we investigated whether Type I IRESs also had the capacity for +1-frame translation. We first surveyed the other dicistrovirus IGR IRESs and their downstream sequences for a potential base pair adjacent to the PKI domain and an overlapping +1 open reading frame both *in silico* (Fig. S1). The IAPV and ABPV IGR IRESs contain an adjacent U-G base pair whereas the SINV-1 and KBV IRESs have a C-G base pair (Fig. S1). All but one of the IGR IRESs contains a potential base pair adjacent to the PKI domain, with the majority of them possessing a potential U-G base pair. As shown previously, the honeybee and fire ant dicistrovirus ORFx proteins are approximately 94-125 amino acids in length (30). The predicted +1 ORFx lengths of other dicistroviruses range from 1-53 amino acids in length. Apart from the honeybee and fire ant dicistrovirus ORFx, the longest putative ORFx sequences are found within the genomes of the CrPV and DCV at 53 amino acids in length.

To determine whether the other IGR IRESs can direct +1-frame translation, we cloned each IGR IRES within the intergenic region of a dual luciferase bicistronic construct (Fig. 1A). The Renilla luciferase (36 kDa; RLuc) monitors scanning-dependent translation whereas firefly luciferase (FLuc) is translated by the IRES. To measure reading frame selection, the firefly luciferase gene is fused in-frame either in the 0 or +1-frame and translated proteins were monitored by incorporation of [^35^S]-methionine/cysteine. Additionally, we generated bicistronic reporter constructs that contain a T2A ‘stop-go’ sequence, which allows quantitation of luciferase activity (32) (Fig. S2). Since we previously showed robust IAPV IRES 0 and +1-frame translation (6), we chose it as a benchmark to compare the +1-frame activity of the other IGR IRESs. In general, all IGR IRESs can direct 0-frame translation to varying extents *in vitro.* Normalizing to the IAPV 0-frame translational activity, the CrPV IGR IRES showed the highest 0-frame translational activity (~170%) whereas the Mud crab dicistrovirus (MCDV) IGR IRES had the lowest activity (~4%; Fig. 1A). By contrast, only a few IGR IRESs can support +1-frame translation. The Type II IGR IRESs from Taura syndrome virus (TSV) and MCDV did not support +1 frame translation suggesting that only a subset of Type II IGR IRESs can direct 0- and +1-frame translation. Interestingly, besides the IAPV IGR IRES, the BQCV and CrPV IRESs mediated +1-frame translation above background levels, 80% and 5% of 0-frame translation, respectively. In summary, only a subset of IGR IRESs can facilitate both 0 and +1-frame translation.

**Fig. 1.**
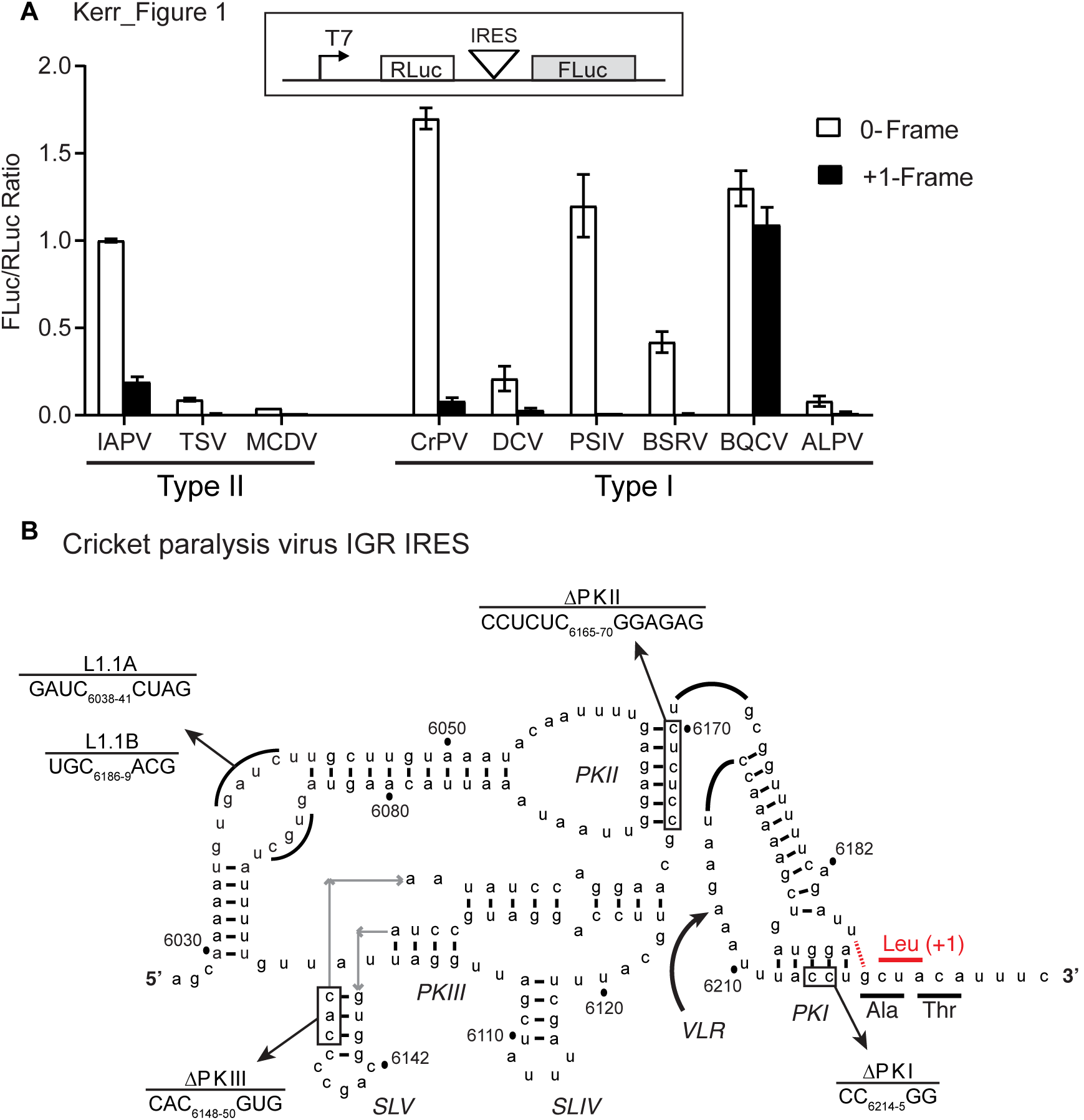
Biochemical analysis of potential +1-frame translation by Dicistrovirus IGR IRESs. A) IGR IRES mediated +1-frame translation using an *in vitro* translation assay. The translation of firefly luciferase (FLuc), which is fused in the 0-frame or +1-frame, is driven by the individual IGR IRES within the bicistronic reporter construct. Linearized reporter constructs were incubated in Sf21 *in vitro* translation insect cell extract at 30 °C for 2 hours in the presence of [^35^S] methionine/cysteine. Translation of FLuc and RLuc was monitored by phosphorimager analysis after resolving on a 16% SDS PAGE. On the bottom panel, the ratio of FLuc/RLuc are quantified and normalized to the IAPV 0-frame translation. Shown are averages from at least three independent biological experiments (± SD). **B) The secondary structure of CrPV IGR IRES**. The ΔPKI, ΔPKII, ΔPKIII, L1.1A, and L1.1B mutants are shown. Comp denotes compensatory mutations combining ΔPKI CC/GG and ΔPKI GG/CC. DM - double mutation of ΔPKI and ΔPKII. TM - triple mutation of ΔPKI, ΔPKII, and ΔPKIII. The potential U6186/G6617 base pair to direct +1 frame translation from a CUA leucine codon.

### CrPV +1-frame translation is IGR IRES-dependent and initiates downstream

To explore +1-frame translation mechanisms further, we focused on the CrPV IRES (Fig. 1B). We first determined whether the structural integrity of the CrPV PKI domain is important for +1-frame translation. For these assays, we used a bicistronic reporter construct that contains the CrPV IRES where a firefly luciferase gene was subcloned into the +1-frame downstream of the IRES with CrPV nucleotides 6217-6387 (Accession KP974707.1), which includes the predicted +1-frame CrPV ORFx (65 kDa; ORFx-FLuc). This dual luciferase reporter construct allows simultaneous monitoring of scanning-dependent and CrPV IRES-mediated 0/+1-frame translation as a shortened 0-frame protein (~11 kD) is also translated in addition to the +1-frame (Fig. 2A). Synthesis of all three proteins is detected by incubating the bicistronic construct in a Sf21 translation extract in the presence of [^35^S]-met/cys (Fig. 2B, lane 1) (6). As shown in Fig. 1A, CrPV IRES +1-frame ORFx translation is approximately 5% of 0-frame translation. Mutating CC_6214-5_ to GG, which disrupts PKI base pairing and abolishes CrPV IRES activity, resulted in negligible or diminished 0- and +1-frame translation whereas a compensatory mutation that restores PKI base pairing rescued translation (10) (Fig. 2B, lanes 2 and 3, Fig. S2), indicating that the integrity of the IRES and the PKI domain is required for CrPV IRES +1-frame translation.

**Fig. 2.**
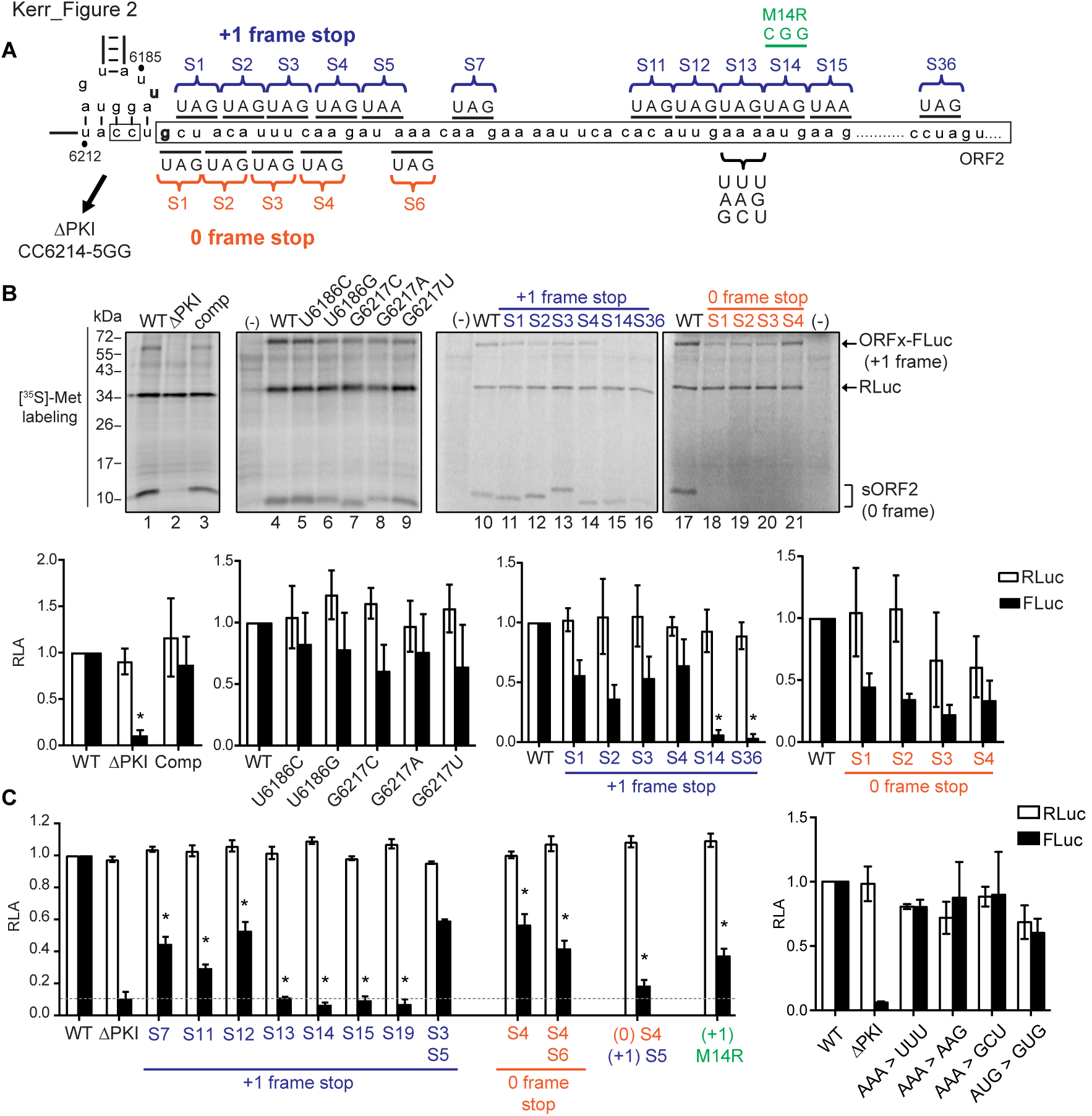
Translation of +1 frame CrPV ORFx is IGR IRES-dependent and initiates downstream of the PKI region. A) Schematic of mutations introduced downstream of the IGR IRES. The PKI region and downstream ‘spacer’ sequence of CrPV IGR IRES are shown. A series of mutations and stop codons were introduced either on the 0-frame or +1-frame. **B) Analysis of the IGR IRES translation *in vitro.*** Linearized reporter constructs are incubated in Sf21 translation extract at 30°C for 2 hours in the present of [^35^S]-methionine/cysteine. Translation of FLuc and RLuc was monitored by autoradiography after resolving on a 16% SDS PAGE. A representative gel from at least three independent biological experiments is shown. * = p-value < 0.05. **C) Quantitation of IRES-mediated translation.** Translation of FLuc and RLuc monitoring FLuc/RLuc enzymatic activity were quantified and normalized to wild-type (WT) IAPV IRES. RLuc monitors scanning-dependent translation acting as an internal control. RLA-relative light units. Shown are averages from at least three independent biological experiments (± SD). * = p-value < 0.05.

The adjacent U-G base pairing of the IAPV IRES is important for +1-frame translation (6). CrPV also has the capacity to form a wobble base pair adjacent to the PKI domain through nucleotides U_6186_ and G_6217_, potentially directing translation from the first +1-frame CUA leucine codon (Fig. 1B). To determine if this base pair is necessary to drive CrPV IRES +1-frame translation, we mutated U_6186_ and G_6217_ to other bases. Mutating U_6186_ to C or G led to an approximate 18%-23% reduction in +1 frame activity and mutating G_6217_ to C, A, or U resulted in roughly a 39%, 24%, and 36% reduction in +1-frame activity, respectively. Although each mutation reduced +1-frame translational activity to some degree, none of the mutations abolished it (Fig. 2B, lanes 4-9). These results suggest that unlike with IAPV, base pairing between nucleotides 6186 and 6217 is not absolutely required for CrPV IRES +1-frame translation.

To determine the potential initiation site of CrPV ORFx, we systematically replaced codons downstream of the IRES with a stop codon and monitored 0- and +1-frame translation *in vitro* using the bicistronic reporter construct (Fig. 2B, 2C). Overall, stop codons placed in the +1-frame did not significantly affect 0-frame translation, indicating that IRES activity was not compromised (Fig. 2B, 2C). Replacing individual codons between the 1st and the 12th +1-frame codon with a stop codon inhibited to varying extents (between 36%-71% reduction compared to wild-type) but did not completely abolish +1 frame translation (Fig. 2B, lanes 11-16; 2C). Conversely, +1-frame translation was completely inhibited when the 13th +1-frame codon and codons thereafter were replaced with a stop codon (Fig. 2B, lanes 15-16; 2C). Replacing both the 3rd and 5th +1-frame codons with stop codons reduced +1-frame translation by 33% but did not eliminate it, suggesting ribosome read-through did not occur. To address the possibility that IRES translation initiates in the 0-frame and then a fraction of translating ribosomes shift into the +1-frame, we inserted a stop codon in the 0-frame downstream of the IRES. As expected, a stop codon in the 0-frame 1st to 4th codons downstream of the IRES abolished 0-frame translation (Fig. 2B, lanes 18-21). However, the 0-frame stop codon insertions reduced by 52-77% but did not eliminate +1 frame translation indicating that ribosomes likely do not shift from the 0 to +1 reading frame. Furthermore, introducing stop codons in both the 0 and +1 frames did not abolish +1-frame translation, though inhibited +1-frame translation by ~80% (Fig. 2C, 0 S4/+1 S5). We noted that the adjacent 14th codon is an AUG methionine. Mutating the AUG to a CGG (M14R) or GUG (M14V) decreased but did not abolish +1-frame translation (Fig. 2C). Mutating the 13th AAA codon to UUU, AAG or GCU did not abolish +1-frame translation (Fig. 2C), suggesting that there is flexibility in the codon identity for CrPV ORFx +1-frame translation. In summary, the mutational analysis indicated that CrPV ORFx translation requires an intact IRES and a +1-frame 13th nonstop codon.

### CrPV +1-frame translation requires 80S assembly and is edeine-insensitive

The current data led us to generate two hypotheses: i) a subset of 40S subunits recruited to the CrPV IRES scan downstream to start translation at the 13th codon (scanning hypothesis) or ii) 40S or 80S ribosomes recruited to the IGR IRES bypass the spacer region to the 13th codon (bypass hypothesis). To address the scanning hypothesis, we took two approaches. First, we utilized mutants of the CrPV IRES in the L1.1 loop that is known to be deficient in recruitment of the 60S subunit (15, 23). If scanning is occurring, 40S subunits recruited to the L1.1 IRES may scan downstream to the downstream +1-frame initiation codon. Reporter constructs harbouring mutations in the L1.1 loop were deficient in 0 and +1-frame translation (Fig. 3A), suggesting that 60S recruitment by the IGR IRES specifically is necessary for translation in the +1-frame. Secondly, we utilized the translational inhibitor edeine to assess if scanning was occurring (Fig. 3B). Edeine prevents the 40S ribosomal subunit from recognizing an AUG start codon (33, 34). Both 0- and +1-frame translation were resistant to edeine relative to that of scanning-dependent translation (Fig. 3B), thus suggesting that 40S scanning is not involved in +1 frame translation.

**Fig. 3.**
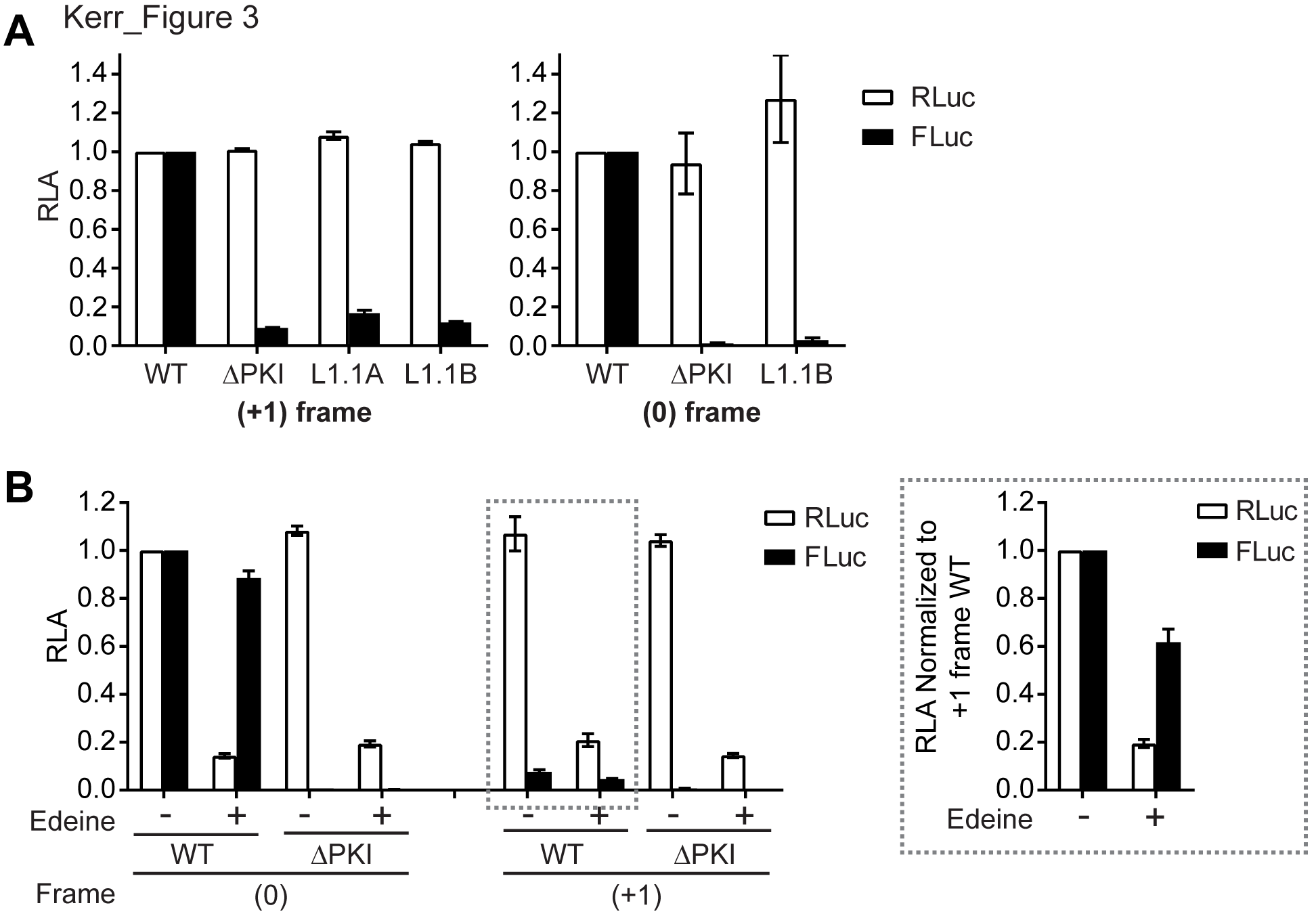
Translation of CrPV ORFx is dependent on 80S binding and scanning-independent. A) Effect of L1.1 mutation on +1 frame translation. Comparison of luciferase activities between WT IGR IRES and L1.1 mutants (L1.1A and L1.1B) that do not bind the 60S subunit. The value of RLuc and FLuc are normalized to WT. **B) +1-frame translation is relatively insensitive to edeine**. CrPV IRES-mediated 0- and +1-frame translation is monitored in the absence or presence of 2 αM edeine. The ΔPKI mutants in both 0- and +1-frame are used as controls for IGR IRES dependent translation. The luciferase activities of RLuc and FLuc on the left panel are normalized to the WT of the 0-frame, whereas the luciferase activities of RLuc and FLuc on the right panel are normalized to the WT of the +1-frame. Shown are averages from at least three independent biological experiments (± SD).

### The integrity of the variable-loop region and pseudo-translocation of the IGR IRES through the ribosome is required for ORFx expression

If 80S ribosomes recruited to the IRES are indeed repositioning or “bypassing” to the downstream 13th +1-frame codon, we next looked to investigate the rules which govern this potential mechanism. Since the IGR IRES, which occupies the ribosomal A site upon ribosome binding, undergoes pseudotranslocation to the ribosomal P site to vacate the A site for delivery of the first aminoacyl-tRNA (17), we reasoned that the ribosome bound to the CrPV IGR IRES must have an empty ribosomal P or A site in order to reposition downstream and accommodate the +1 frame start codon. To address this, we introduced mutations in the variable loop region (VLR), which has been shown to disrupt the IGR IRES-mediated pseudotranslocation event (Fig. 1B) (35). Specifically, shortening the length of the VLR by 2 or 3 nucleotides (Δ2 and Δ3, respectively) inhibits the first pseudo-translocation event from the A site to the P site whereas altering the identity of nucleotides A_6204_, and AA_6208-9_ to guanosines (G-rich) inhibited IRES translocation from the P site to the E site (35). Interestingly, all three VLR mutants decreased +1-frame activity (Fig. 4); both the G-rich and Δ3 mutants demonstrated little to no activity, while the Δ2 mutant still exhibited roughly 50% activity to that of WT. This result is consistent with previous data that the Δ2 mutant IRESs are still able to accommodate a fraction of aminoacyl-tRNA in the A site (~25%), allowing translation to occur (35). Altogether, these results indicate that the pseudotranslocation event of the IGR IRES through the ribosome contributes to +1-frame translation downstream.

**Fig. 4.**
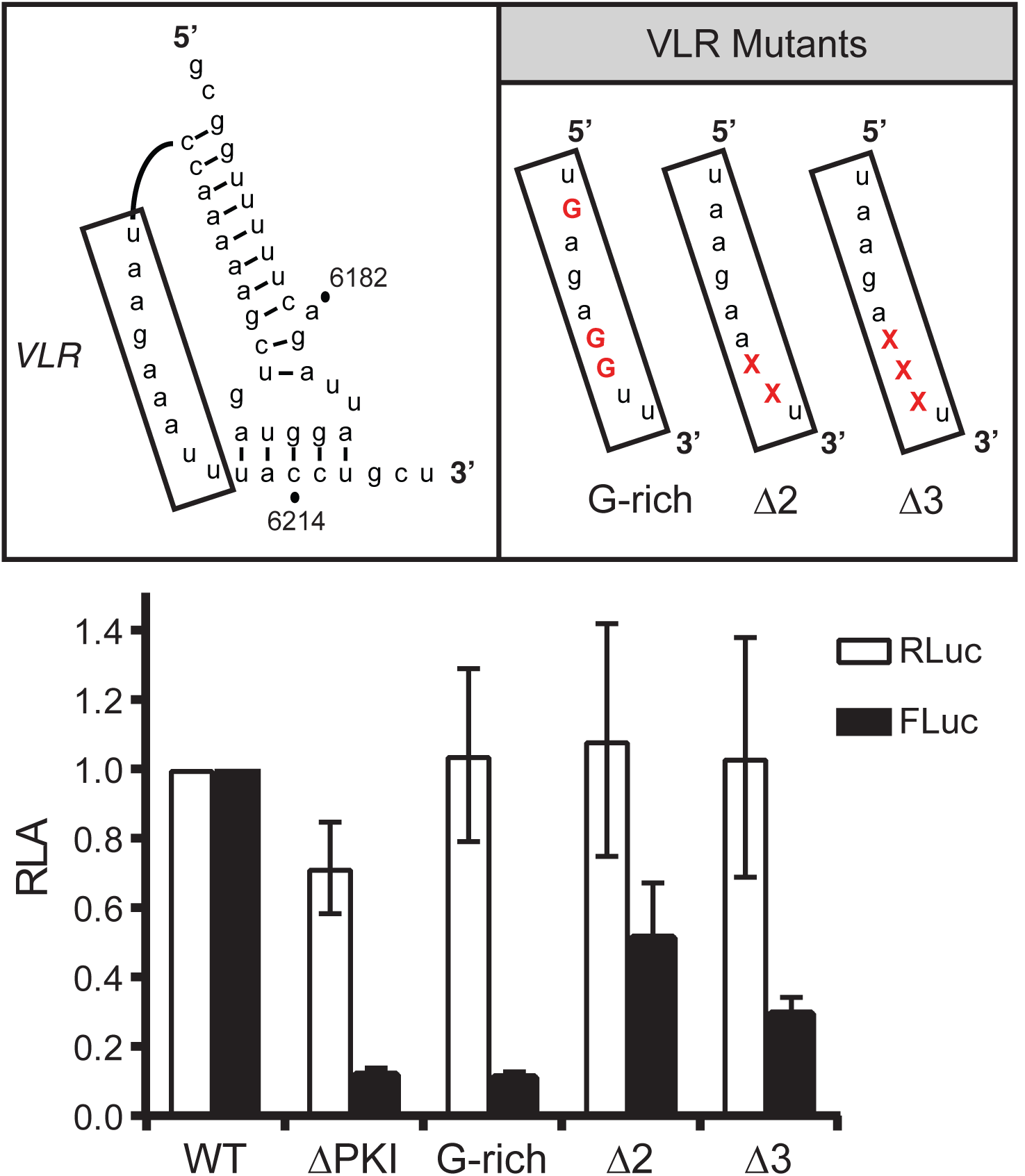
Pseudotranslocation is necessary for +1-frame translation. Constructs containing mutations in the variable loop region (VLR) of the IGR IRES known to disrupt pseudotranslocation were monitored for translation activity. Luciferase activities were compared between WT IGR IRES and VLR mutants after *in vitro* translation assays. “x” denotes deleted nucleotides within VLR. Shown are averages from at least three independent biological experiments (± SD).

### The spacer region downstream of the IGR IRES is necessary for +1 frame translation

We next investigated whether the spacer region located between PKI of the IGR IRES and the downstream AAA codon contributes to IRES-mediated ribosome bypass. Our results showed that inserting stop codons in the ‘spacer region’ (Fig. 2A) between the IRES PKI domain and the +1-frame 13th codon of ORFx inhibited but did not completely abolish +1-frame translation, suggesting that an element within this spacer region may promote +1-frame translation. We first addressed whether the CrPV spacer region is sufficient to direct +1-frame translation by generating a chimeric construct whereby the PSIV IRES is fused with the CrPV spacer region (Fig. 5). The PSIV IRES directs strong 0-frame but no or relatively weak +1-frame translation (Fig. 1A). The PSIV-CrPV spacer chimeric reporter resulted in 0-frame translation (Fig. 5A), indicating that the spacer region does not affect the activity of the PSIV IRES. In contrast to the PSIV IRES construct, the chimeric PSIV-CrPV reporter resulted in +1-frame translation, implying that the CrPV spacer region is sufficient to drive +1-frame translation in the presence of a functional IGR IRES (Fig. 5A).

**Fig. 5.**
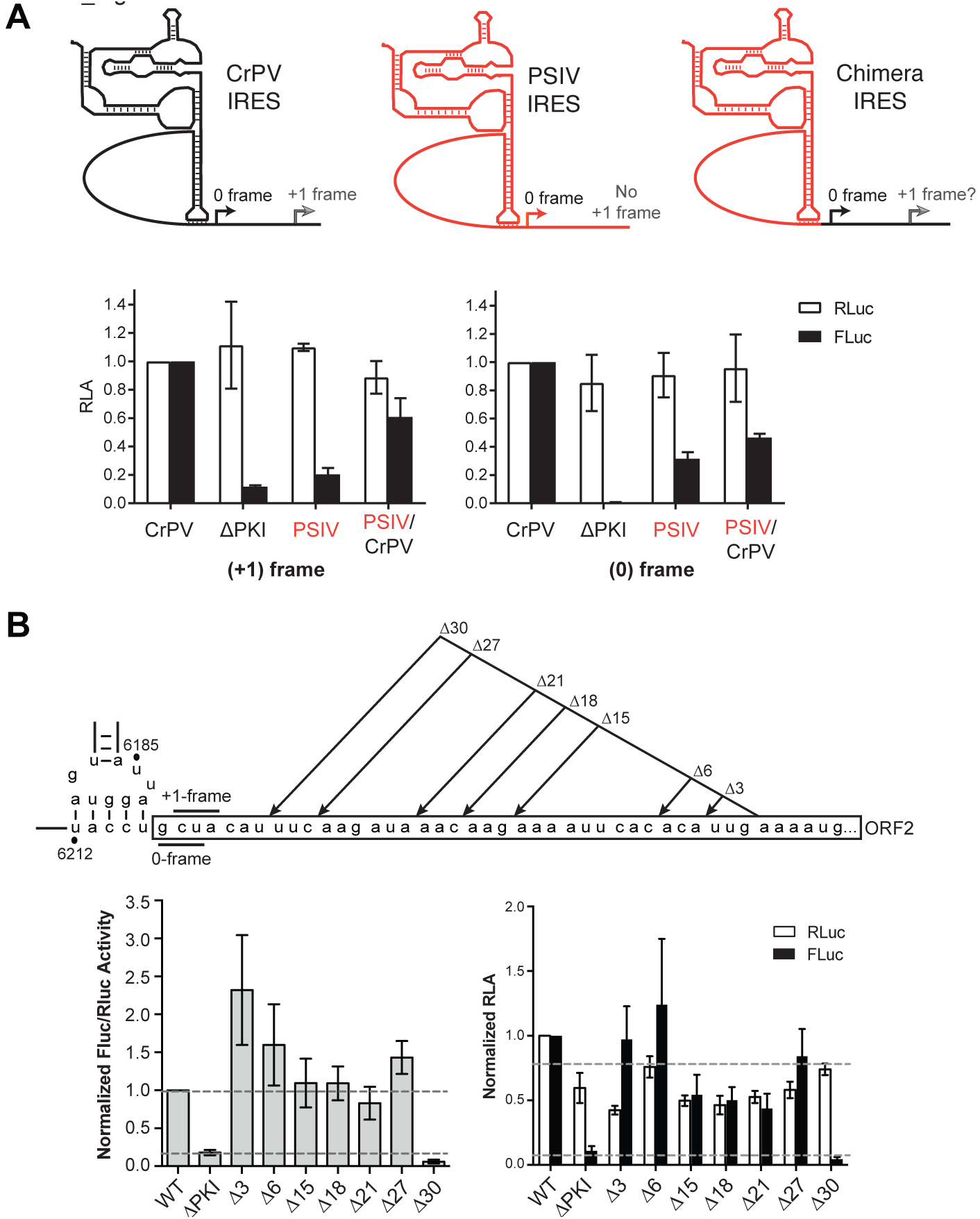
The downstream sequence of CrPV IGR IRES promotes +1-frame translation. A) Chimeric IRES displays +1-frame translation activity. The PSIV IGR IRES, which shows no +1-frame translation activity (Fig. 1 and Fig. 3A), is combined with the downstream sequence of CrPV IGR IRES PKI region. The luciferase activities are normalized to the wild-type CrPV constructs in both +1- and 0-frame. The PSIV-CrPV chimera in the 0-frame is used as control to demonstrate that PSIV IGR IRES functions under this context. B) Truncation of the spacer region impairs +1-frame translation from the CrPV IGR IRES. Schematic displaying truncations within the spacer region between the PKI domain and the +1 frame start site is shown above. *In vitro* translation assay of the CrPV IRES bicistronic construct with the respective mutations were monitored by luciferase activities. Both RLuc and FLuc activities are normalized to that of WT constructs. Results are shown as a normalized ratio of FLuc/RLuc activity (left) and as separated RLuc and Fluc activities (right) Shown are averages from at least three independent biological experiments (± SD). RLA = relative luciferase activity

To delineate whether there is a specific element within the spacer region that is required for CrPV +1 frame translation, we systematically deleted from the 3’ end of the spacer region (Fig. 5B). Interestingly, deleting 3 to 27 nucleotides did not affect +1 frame translation. However, deleting 30 nucleotides and leaving seven nucleotides adjacent to the IRES abolished +1 translation (Fig. 5B). Δ3, Δ6 and Δ27 mutants appear to have much higher Fluc/Rluc activity when compared to WT, an observation that requires further examination. In summary, these results suggest that the specific sequences and its context within the spacer region, particularly the sequences immediately downstream of the PKI domain, are important for mediating IRES-dependent +1 frame translation.

### +2-frame translation mediated by the CrPV IRES

We hypothesized that if the ribosome is indeed bypassing to a specific ‘landing site’ then it should be independent of frame, whereas if the ribosome merely begins translation in the 0-frame before slipping into the +1-frame, then we expect to see no ORFx translation. To address this, we inserted a series of nucleotides into our bicistronic construct that shifts only the ORFx-Fluc into the +2-frame (Figure S3). Specifically, we inserted either a single nucleotide or up to 7 proceeding the 13^th^ AAA codon. As a control, we inserted either 6 or 9 nucleotides in the same position that does not introduce an additional frameshift. Insertion of a U creates a stop codon in the +1-frame while an inserted C does not. To our surprise, we observed ORFx expression with all insertions (Figure S3). These results suggest translation of ORFx is reading frame-independent and indicates that the ribosome may be repositioned from the IRES to the downstream 13^th^ codon.

### CrPV +1-frame ORFx is expressed yet not required for infection in *Drosophila* S2 cells

Our *in silico* and biochemical data indicate that ribosomes recruited to the CrPV IRES may bypass downstream to the +1-frame translation from the 13th codon. The CrPV ORFx is predicted to be 41 amino acids in length if ORFx is translated from the +1-frame 13th codon (Fig. 6). To determine whether CrPV ORFx is synthesized during infection, *Drosophila* S2 cells infected with CrPV (MOI 10) were harvested at 6 hours post infection and lysed. Proteins were subsequently digested with trypsin and peptides were analyzed by LC-MS/MS. We identified two peptides that correspond to CrPV ORFx both of which were located downstream of the +1-frame 13^th^ codon (Fig. 6). Importantly, these peptides were not identified in mock-infected S2 cells. Thus, ORFx is expressed in CrPV-infected S2 cells.

**Fig. 6.**
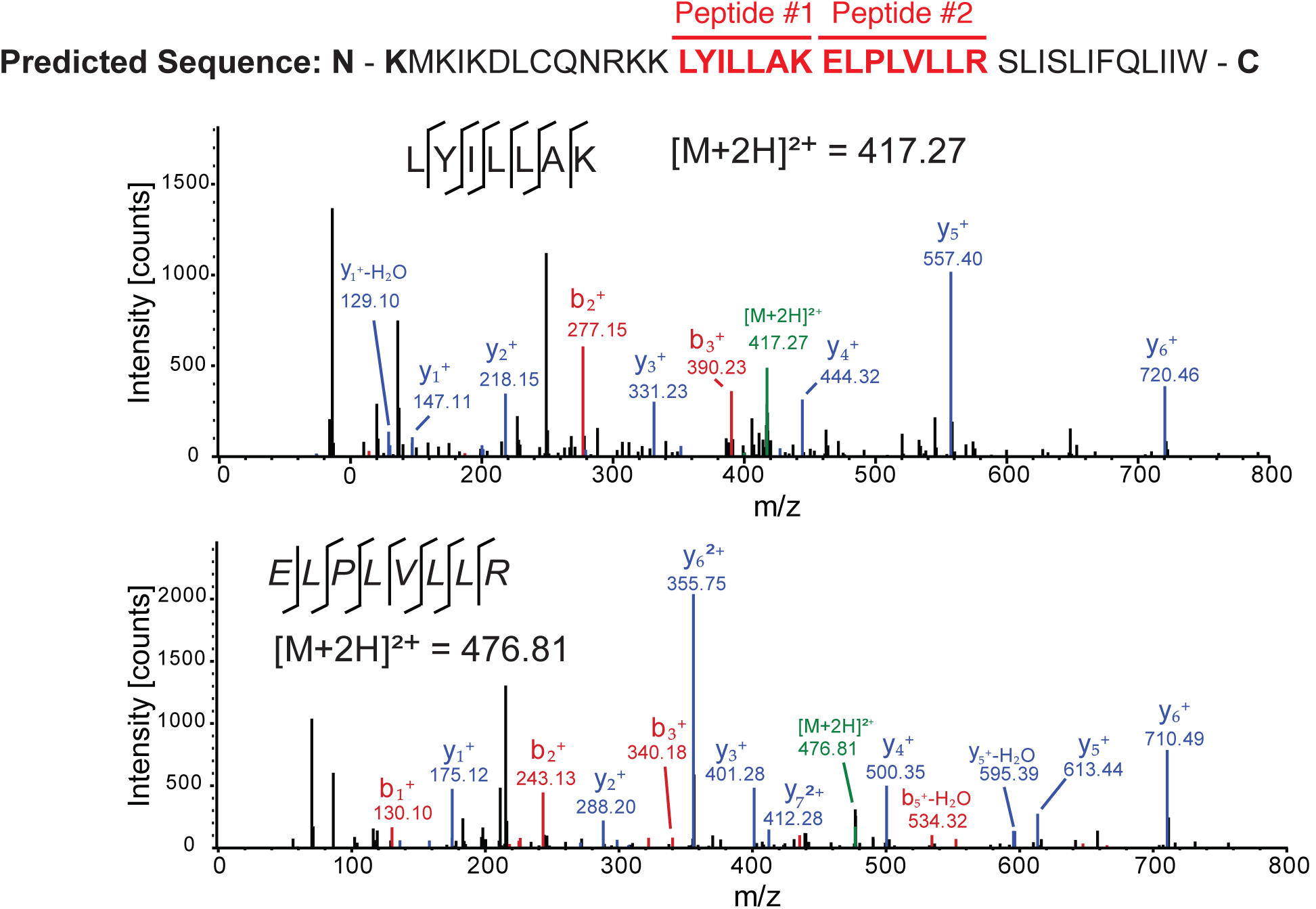
ORFx is expressed in CrPV infected S2 cells. The predicted +1 frame amino acid sequence is shown above. Residues that are italicized represent the amino acid sequence if initiation occurred adjacent to the IGR IRES, whereas the bolded K lysine residue denotes the position of the 13^th^ amino acid. *Drosophila* S2 cells were infected with CrPV (MOI 10). Proteins were extracted, digested with trypsin and subjected to LC-MS/MS analysis. Peptide fragment spectra were searched against a *Drosophila* uniprot database plus CrPV proteins. Two peptides (highlighted) were detected from the trypsin digestion of S2 cell lysate at 6 hours post infection. Individual fragment ions are annotated in the spectra and in the sequence representation.

Given this, we sought to determine the influence of CrPV ORFx expression on the outcome of viral infection. Using a recently developed CrPV infectious clone, termed CrPV-2, we introduced mutations that would abolish ORFx expression (36). To this end, we created two separate mutant clones: the +1-frame 12^th^ codon (UUG_6251-3_) was altered to an amber stop codon (UAG; CrPV-S12) and the +1-frame 19^th^ codon (UUA_6272-4_) changed to an ochre stop codon (UAA; CrPV-S19) (Fig. 7A). Both mutations are synonymous in the 0-frame. Based on our translation data (Fig. 2), the +1-frame S19 but not the S12 would inhibit ORFx expression.

**Fig. 7.**
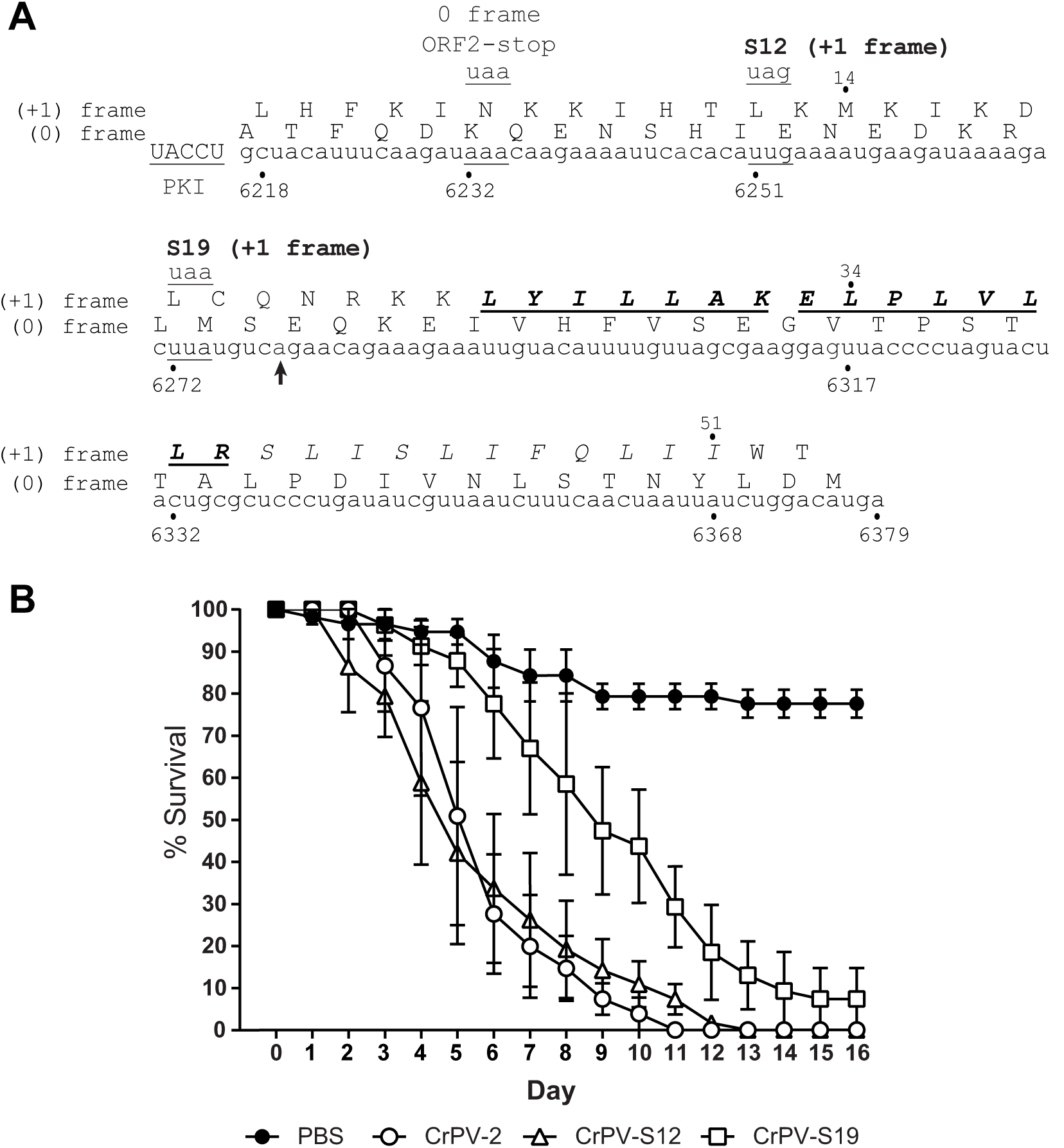
Pathogenesis of a CrPV mutant lacking ORFx is attenuated in adult *Drosophila melanogaster* flies. A) Schematic of mutations in the CrPV infectious clone, CrPV-2. The downstream nucleotide sequence of CrPV-2 IGR IRES and its potential amino acid sequence of +1-frame ORFx are shown. The ORF2-Stop mutant depletes the synthesis of 0-frame viral structural proteins. Mutants S12 and S19 place a stop codon on the +1-frame with a synonymous mutation in the 0-frame. Note that CrPV-2 (Accession: KP974706) sequence has a C_6279_A mutation (denoted with a black arrowhead) compared to the original CrPV (Accession: AF218039) sequence. Residues that are italicized, underlined and in bold are the peptides identified by mass spectrometry in Fig. 6. **B)** Adult flies (Iso w^1118^; 10 male, 10 female) were injected intrathoracically with 5000 FFU of CrPV-2, CrPV-S12, CrPV-S19, or PBS. Subsequently, flies were flipped onto standard media and survival was monitored daily. Shown is a graph representing the average from three separate biological experiments (±SEM).

First, we examined whether ORFx influences viral protein synthesis *in vitro.* Incubation of *in vitro* transcribed CrPV-2 RNA in Sf21 translation extracts led to synthesis and processing of all viral proteins as reported previously (Fig. S4A) (36). While stop codons in the 0-frame of ORF1 or ORF2 inhibited synthesis of viral proteins, both CrPV-S12 and CrPV-S19 RNAs resulted in viral protein synthesis that was indistinguishable compared to wild-type CrPV -2 *in vitro,* demonstrating that CrPV ORFx is not necessary for viral protein synthesis *in vitro* (Fig. S4B) (36).

We next assessed the viability of the CrPV-S12 and -S19 viruses in cell culture. Harvested wild-type, mutant S12 or S19 CrPV-2 virus were used to infect naïve S2 cells at a MOI 10 and 1 to follow the first round of infection and subsequent rounds of infection, respectively. Infection with wild-type CrPV-2, CrPV-S12 or CrPV-S19 all resulted in accumulation of viral proteins and RNA and shutdown of host translation in a similar manner (Fig. S4C). Similarly, neither infection produced significantly different titres between wild type CrPV-2 and either CrPV-S12 or CrPV-S19 at any time point, apart from both mutant viruses resulting in higher titres than CrPV-2 after 24 hours post infection (Fig. S4D). A similar result was observed with infections at a MOI 0.1 (data not shown). Taken together, ORFx has no observable effect on the life cycle of CrPV in S2 cells.

### ORFx contributes to CrPV infection in adult flies and associates with membranes

We addressed whether ORFx contributes to CrPV infection in adult fruit flies. To test this, we injected adult flies intrathoracically with PBS, CrPV-2, CrPV-S12, or CrPV-S19 and monitored fly mortality daily. Flies injected with CrPV-2 or CrPV-S12 exhibited 50% mortality by day 5 and 100% mortality by day 11 and 12 (Fig. 7B). By contrast, flies injected with CrPV-S19 did not reach 50% mortality until day 9 and reached 100% mortality at day 14 (Fig. 7B). These results demonstrate that ORFx contributes to CrPV pathogenesis in adult flies.

To determine if the effect seen on CrPV pathogenesis is a result of defects in viral replication, we measured viral titres and assessed viral protein levels in injected adult flies at 5 days post infection. Interestingly, viral titres and viral protein levels showed no significant differences between wild type CrPV-2, CrPV-S12, and CrPV-S19 (Fig. S5). Finally, using RT-PCR followed by sequencing, the S12 and S19 mutations are stable during virus propagation in S2 cells (data not shown). In summary, our results indicate that the defect in viral pathogenesis observed in the CrPV-S19 mutant virus is not due to a defect in viral replication.

From our data, CrPV ORFx is predicted to be a 41 amino acid protein, mediated by a ribosomes bypass mechanism from the 13th +1-frame AAA codon. Comparing the sequence of ORFx from CrPV and other species shows no appreciable homology to other proteins. Nevertheless, *in silico* topology predictions suggest that ORFx can adopt an alpha helical transmembrane segment at its C-terminus (amino acids 22-39; Fig. S6A). To examine ORFx function, we generated constructs containing either N- or C-terminal HA-tagged ORFx. Transfection for 10 or 24 hours with the HA-tagged ORFx constructs did not result in a dramatic decrease in S2 cell viability as measured by a trypan blue exclusion assay (Fig. S6B). However, by 48 hours, transfection of ORFx-HA and HA-ORFx led to a relatively minor reduction (13% and 15% decrease, respectively) in cell viability, suggesting that ORFx expression is slightly toxic in S2 cells. Immunoblotting for HA showed that HA-ORFx is expressed in S2 cells (Fig. S6C). To determine the subcellular localization in S2 cells, we transfected the HA-tagged ORFx constructs and monitored ORFx localization (pORFx-HA) by HA-antibody immunofluorescence staining in comparison with cytoplasmic, ER, Golgi, and nuclear marker protein antibodies. We also mutated two pairs of amino acids, LV and LI, to KK and KR respectively, to disrupt the transmembrane domain (pORFx-TMmut-HA). Co-staining showed that the wild-type HA-tagged ORFx overlaps mainly with ER protein marker Calnexin, and partially overlaps with Golgi-associated protein, Golgin84 (Fig. 8, S7, pORFx-HA). The wild-type HA-ORFx displayed little to no overlap with α-Tubulin staining. By contrast, the mutant transmembrane HA-tagged ORFx staining showed no overlap with Calnexin or Golgin84 (Fig. 8, S7, pORFx-TmMut-HA). Collectively, these results suggest that ORFx associates with membranous organelles and potentially the ER specifically. To examine this further, we used differential centrifugation to separate subcellular components followed by immunoblotting of HA. As expected, tubulin is found in the cytoplasmic fraction and the ER-associated KDEL protein and cytochrome C are enriched in nuclei and ER fractions (Fig. S6C). Both HA-tagged ORFx are detected within membranous fractions but not within the cytoplasmic fractions (Fig. S6C), further supporting that ORFx resides within the membrane of cells that may be important for its function.

**Fig. 8.**
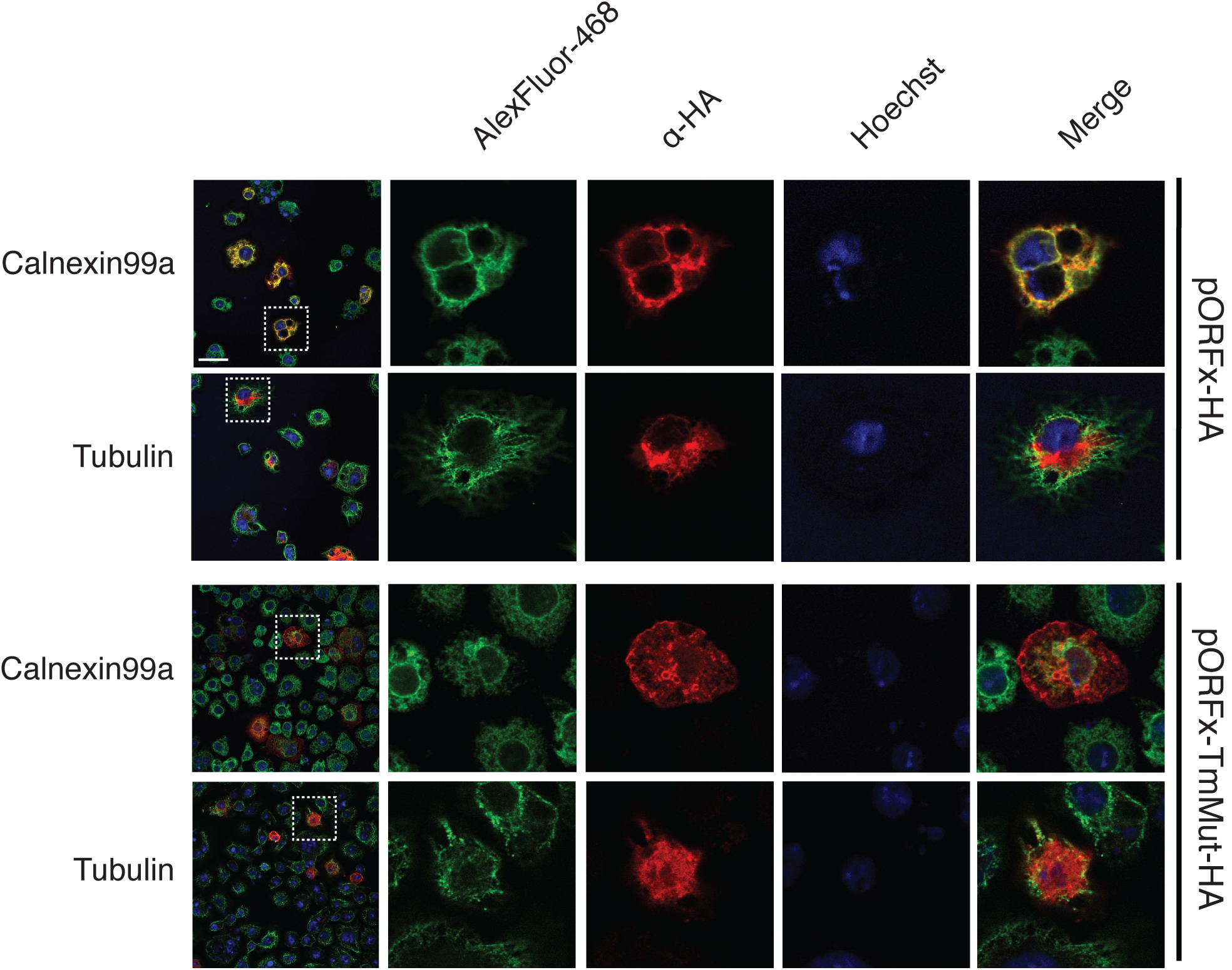
Subcellular localization of CrPV ORFx. S2 cells were transfected with a construct expressing C-terminally HA-tagged ORFx or a mutant version of ORFx in the transmembrane region and incubated for 48 h. Following incubation, cells were fixed, permeabilized and co-stained with HA antibody and antibodies against the ER (Calnexin 99A) and cytoplasm (alpha-Tubulin). Shown are representative Z-stack micrographs from three independent experiments. Scale bars represent 15 μm.

## Discussion

Recoding mechanisms have illuminated diverse RNA structural elements that interact with the ribosome to affect reading frame maintenance (2). In this study, we have demonstrated a novel translation recoding mechanism by which an IRES may promote ribosome repositioning at a downstream codon. In contrast to the IAPV IGR IRES, which directs ribosome reading frame selection (6), the CrPV IGR IRES can facilitate the expression of a downstream +1 overlapping frame, which we termed ORFx. We provide extensive analysis using mutagenesis that translation of CrPV ORFx likely occurs 37 nucleotides downstream at the 13^th^ AAA (Lys) codon. Moreover, we show that ORFx is expressed in CrPV-infected cells by mass spectrometry and that ORFx is required for promoting CrPV pathogenesis in a *Drosophila* injection model. Our data suggest a model whereby after 80S assembly on the CrPV IGR IRES, the majority of the ribosomes translate in the 0-frame by delivery of the incoming Ala-tRNA^Ala^ to the 0-frame GCU codon whereas a fraction of ribosomes bypasses or “slides” 37 nucleotides downstream to direct translation at the +1-frame AAA (Lys) codon (Fig. S8). From our results, the following rules appear to apply to +1-frame translation in CrPV: i) the PKI must be intact, ii) both 40S and 60S ribosomal subunits are required to bind to the IRES, iii) pseudotranslocation of the IRES through the ribosome is necessary, iv) the spacer region between PKI and the 13^th^ AAA codon is essential, and v) the nucleotide identity of the spacer region is crucial for efficient +1 frame translation.

A unique feature of this repositioning mechanism is that an intact IRES is essential for +1-frame translation (Fig. 2). The current model is that the PKI domain of the IRES occupies the ribosomal A site upon ribosome binding to the IRES, followed by a pseudotranslocation event in order to vacate the A site to allow delivery of the first aminoacyl-tRNA in the 0-frame (17). Given this model, it is difficult to envision how the IRES can direct the ribosome to bypass 37 nucleotides to initiate translation at a +1-frame AAA Lys codon. A potential clue comes from our mutagenesis analysis that suggests that translocation of the CrPV IRES through the ribosome is a prerequisite for downstream +1-frame translation (Fig. 4). This is intuitive as the PKI domain must vacate the A site in order to allow translation in both the 0 and +1 frames. At this point, it is unclear whether the CrPV IGR IRES ribosome repositioning event requires delivery of an aminoacyl-tRNA to the A site of the ribosome prior to the ribosome sliding downstream to initiate translation at the 13th +1 frame codon or whether a vacant ribosome bypasses from the IRES to the initiating +1 frame codon. Determination of the aminoacyl-tRNA(s) that are delivered during the initial CrPV IGR IRES pseudotranslocation event should provide insights whether the ribosomal A site is vacant prior to ribosome sliding downstream to the 13th +1 frame codon.

A recent *in vitro* study using single molecule fluorescence spectroscopy demonstrated that the CrPV IGR IRES can facilitate +1-frame translation approximately 5% of the time compared to 0-frame translation whereby reading frame selection is dictated by the kinetics of tRNA binding in the first 0- or +1-frame codon (12). Using a more physiologically-related system, our results suggest that the CrPV IRES directs +1-frame translation using a different mechanism. Based on our systematic stop codon mutational analyses, the most likely scenario is that ribosomes recruited to the IRES must bypass downstream to translate in the +1-frame (Fig. 2). In support of this, insertion of a stop codon at the 12th +1-frame codon, but not the 19th, attenuates CrPV-mediated death in a Drosophila injection model, thus providing biological significance of CrPV +1-frame translation (Fig. 7). We also note that the downstream authentic ‘spacer’ sequence is necessary for +1-frame translation (Fig. 5), which is partially absent and/or altered from previous studies, which may preclude detection of bypass (12).

The downstream spacer region appears to be a key feature that is necessary for efficient CrPV IRES-mediated +1-frame synthesis (Fig. 5). Interestingly, the majority of mutations within the spacer region causes a reduction ranging from 10-80% in the amount of CrPV IRES-mediated +1-frame synthesis (Fig. 2), suggesting that nucleotide or codon identity is crucial for triggering the bypass event. There are no obvious RNA secondary structures within the spacer region, however, we cannot rule out any long distance RNA:RNA interactions that may contribute to +1-frame translation.

Furthermore, sequences immediately downstream of the IRES PKI domain are important for +1 frame translation (Fig. 5B). Investigation into how the spacer sequence influences ribosomes bound to the IRES to start translation in the 0-frame or at the downstream +1-frame initiating codon should shed light onto this atyptical ribosome bypass mechanism.

How does the ribosome reposition to the downstream +1-frame initiation codon after IRES binding? It is possible that the repositioning of the ribosomes occurs via mechanism similar to that observed with prokaryotic 70S ‘scanning’ (37) or 70S ‘sliding’ that occurs in coupled translational reinitiation (38, 39) (Fig. S8). Indeed, a related dicistrovirus PSIV IRES can direct translation using a scanning-like mechanism in prokaryotes, which may suggest a similar property observed with our studies on the CrPV IRES (40). Moreover, it has been reported and proposed that energy-independent scanning or diffusion of the ribosome or ribosomal subunits (i.e. phaseless wandering) can occur to locate an AUG codon (39, 41-43)(Fig. S8). Nevertheless, our study shows that CrPV IRES +1-frame translation requires 80S ribosome binding to the IRES (Fig. 3A) and is edeine-insensitive (Fig. 3B), thus we favour a model that 80S ribosomes reposition to the 13th +1-frame codon. This warrants comparison with the translational bypassing observed in *gene 60* of T4 bacteriophage (44). In *gene 60,* translating ribosomes stall in a non-canonical rotated state at a ‘take-off’ Gly codon with a peptidyl-tRNA^Gly^, which dissociates from the anticodon, and ‘lands’ at a matching Gly codon 50 nucleotides downstream and allowing translation to resume. This bypass mechanism is dependent on a post-translocation step requiring a downstream 5’ ‘take-off’ stem-loop structure and a nascent translated peptide (45, 46). Although CrPV does not have an obvious RNA structure within the spacer region, it is possible that the highly complex structure of the IGR IRES itself may contribute to downstream +1 frame translation, especially given its dynamic nature during movement through the ribosome (Fig. S8)(28, 47). Moreover, it is known that the ribosome bound to the IGR IRES is in a rotated state and that the first pseudotranslocation step is rate limiting, which may contribute to CrPV IRES reading frame selection (17, 29). Finally, it is possible is that CrPV IRES bypass could be occurring through an RNA looping event (48); the downstream RNA is brought into close proximity with the 80S ribosome allowing it to transition to 13^th^ AAA codon potentially by an unknown protein factor or a long-range RNA:RNA interaction. Taken together, it is likely that it is a combination of tRNA kinetics and conformational changes of the IRES and the downstream spacer region that lead to bypassing, of which the contributions of each element require further investigation.

The biological relevance of CrPV +1-frame translation was initially evidenced by the detection of ORFx peptides in CrPV-infected S2 cells (Fig. 6). To our surprise, disruption of ORFx synthesis by stop codon insertion in the +1-frame did not perturb viral infection in tissue culture cells but showed retarded mortality in adult flies even though viral load remained similar between wild type and mutant viruses (Fig. 7). CrPV infection is thought to cause death through paralysis, subsequently leading to dehydration or starvation of the host (49, 50). CrPV can infect several tissues in the fly including the trachea, midgut, and central nervous system although the latter has not been demonstrated directly (48, 50, 51). How ORFx may contribute to CrPV pathogenesis is an outstanding question. Our results indicate that ORFx is membrane associated (Fig. 8, S7) and does not contribute directly to viral replication but rather to the pathogenesis of CrPV infection in fruit flies (Fig. 7, S4). Furthermore, expression of ORFx is slightly cytotoxic in Drosophila cells, a property that may also contribute to pathogenesis of CrPV infection. Future studies into the localization of ORFx, potential interacting partners, and tissue tropism of wild-type versus mutant virus infection in the fly should provide insights into the role of ORFx.

Viruses continue to surprise us with their ability to manipulate the ribosome in remarkable ways. Here, in addition to the previous findings on the honey bee dicistrovirus IRES, we have revealed another recoding mechanism utilizing an IRES, thus highlighting the strong selection to increase the coding capacity of the dicistrovirus genome. Furthermore, an IRES that can direct ribosome repositioning to facilitate the translation of a hidden +1 overlapping ORFx adds to the growing list of diverse pathways of ribosome translational recoding. It will be of considerable interest to investigate whether other IRESs direct reading frame selection by a similar CrPV IGR IRES-mediated ribosome repositioning mechanism. Ribosome repositioning or bypass is not specific to bacteria or mitochondria (52, 53) but may be a more general phenomenon in eukaryotes that initially thought.

## Materials and Methods

Cell culture and virus. *Drosophila* Schneider line 2 (S2; Invitrogen) cells were maintained and passaged in Shield’s and Sang medium (Sigma) supplemented with 10% fetal bovine serum.

Propagation of CrPV in *Drosophila* S2 cells has been previously described (54). CrPV-2 and mutant viruses were generated from *Drosophila* S2 cells using an adapted protocol (55). Briefly, 5.0 ×10^7^ S2 cells were transfected with *in vitro* transcribed RNA derived from pCrPV-3 or mutant plasmids and incubated for 48h. Cells were dislodged into the media, treated with 0.5% Igepal CA-630 (Nonidet P-40) and 0.1% 2-mercaptoethanol, and incubated on ice for 10 min. Cell debris was cleared by centrifugation at 13,800 RCF for 15 min at 4°C. Viral particles were then concentrated by ultracentrifugation at 141,000 RCF for 2.5 h at 4°C. The pellet was resuspended in PBS and sterilized through a 0.2 μM filter. Viral titres and yield were determined as previously described (20). All viruses were sequence verified via RT-PCR with primers directed against the CrPV IGR IRES.

LC-MS/MS Analysis. Cell pellets harvested from CrPV-infected cells at 6 hpi were solubilized in 1% sodium deoxy cholate and 50 mM NH_4_HCO_3_. Protein concentrations were determined via BCA assay (Thermo). Proteins (100 μg) were reduced (2 μg DTT, 37°C, 30 min) and alkylated (5 μg iodoacetamide, RT, 20 min). Samples were digested with trypsin overnight at room temperature. Peptides were acidified with 1% TFA to pH <2.5 and the precipitated deoxycholate was remove via centrifugation. Peptides were desalted and concentrated on C18 STAGE-tips, eluted in 80% acetonitrile, 0.5% acetic acid, and dried in a vacuum concentrator (Eppendorf)(Rappsilber, Ishihama, & Mann, 2003). Samples were resuspended in 20% acetonitrile and 0.1% formic acid before loading on an Agilent 6550 mass spectrometer.

Data were searched using MaxQuant (v1.5.3.30)(56). Parameters included: carbamidomethylated cysteine (fixed), methionine oxidation (variable), glutamine and asparagine deamidation (variable), and protein N-terminal acetylation (variable); trypsin specific; maximum 2 missed cleavages; 10 ppm precursor mass tolerance; 0.05 Da fragment mass tolerance; 1% FDR; +1 to +7 charge states; and common contaminants were included. Both the *Drosophila* and CrPV protein databases used were the most recent annotations downloaded from UniProt (www.uniprot.org).

Fly stocks and viral injections. Flies *(Isogenic w^1118^;* Bloomington Drosophila Stock Center) were maintained on standard cornmeal food at 25°C and 70% humidity with a 12 h light-dark cycle. Freshly eclosed virgin males and females were separated and collected in groups of 10 each. Flies (10 males and 10 females) were injected with 200 nL of PBS, CrPV-2, CrPV-S12, or CrPV-S19 (5000 FFU) using a PV830 PicoPump (World Precision Instruments) and transferred to standard food. Mortality was monitored daily.

## Acknowledgements

We thank John Atkins and the Jan lab for insightful discussions. Kathy Pan and Seonghoon Lee contributed a subset of data in Fig. 1A and 2B, respectively. This study was supported by a Canadian Institutes of Health Research (PJT-148761) to E.J. and to D.A. (MOP-130517), an NSERC Discovery Grant to L.F. and an NSERC CGS-D fellowship to C.K.

## References

1 Baranov PV, Atkins JF, & Yordanova MM (2015) Augmented genetic decoding: global, local and temporal alterations of decoding processes and codon meaning. Nature reviews. Genetics 16(9):517–529.

2 Dinman JD (2012) Mechanisms and implications of programmed translational frameshifting. Wiley interdisciplinary reviews. RNA 3(5):661–673.

3 Firth AE & Brierley I (2012) Non-canonical translation in RNA viruses. J Gen Virol 93(Pt 7):1385–1409.

4 Jan E, Mohr I, & Walsh D (2016) A Cap-to-Tail Guide to mRNA Translation Strategies in Virus-Infected Cells. Annual review of virology 3(1):283–307.

5 Au HH & Jan E (2014) Novel viral translation strategies. Wiley interdisciplinary reviews. RNA 5(6):779–801.

6 Ren Q, et al. (2012) Alternative reading frame selection mediated by a tRNA-like domain of an internal ribosome entry site. Proc Natl Acad Sci U S A 109(11):E630–639.

7 Jackson RJ, Hellen CU, & Pestova TV (2010) The mechanism of eukaryotic translation initiation and principles of its regulation. Nature reviews. Molecular cell biology 11(2): 113–127.

8 Wilson JE, Pestova TV, Hellen CU, & Sarnow P (2000) Initiation of protein synthesis from the A site of the ribosome. Cell 102(4):511–520.

9 Jan E, Kinzy TG, & Sarnow P (2003) Divergent tRNA-like element supports initiation, elongation, and termination of protein biosynthesis. Proc Natl Acad Sci U S A 100(26):15410–15415.

10 Jan E & Sarnow P (2002) Factorless ribosome assembly on the internal ribosome entry site of cricket paralysis virus. J Mol Biol 324(5):889–902.

11 Pestova TV & Hellen CU (2003) Translation elongation after assembly of ribosomes on the Cricket paralysis virus internal ribosomal entry site without initiation factors or initiator tRNA. Genes Dev 17(2):181–186.

12 Petrov A, Grosely R, Chen J, O’Leary SE, & Puglisi JD (2016) Multiple Parallel Pathways of Translation Initiation on the CrPV IRES. Mol Cell 62(1):92–103.

13 Nishiyama T, et al. (2003) Structural elements in the internal ribosome entry site of Plautia stali intestine virus responsible for binding with ribosomes. Nucleic Acids Res 31(9):2434–2442.

14 Costantino DA, Pfingsten JS, Rambo RP, & Kieft JS (2008) tRNA-mRNA mimicry drives translation initiation from a viral IRES. Nat Struct Mol Biol 15(1):57–64.

15 Pfingsten JS, Costantino DA, & Kieft JS (2006) Structural basis for ribosome recruitment and manipulation by a viral IRES RNA. Science 314(5804):1450–1454.

16 Schuler M, et al. (2006) Structure of the ribosome-bound cricket paralysis virus IRES RNA. Nat Struct Mol Biol 13(12):1092–1096.

17 Fernandez IS, Bai XC, Murshudov G, Scheres SH, & Ramakrishnan V (2014) Initiation of Translation by Cricket Paralysis Virus IRES Requires Its Translocation in the Ribosome. Cell.

18 Au HH, et al. (2015) Global shape mimicry of tRNA within a viral internal ribosome entry site mediates translational reading frame selection. Proc Natl Acad Sci U S A 112(47):E6446–6455.

19 Muhs M, et al. (2015) Cryo-EM of ribosomal 80S complexes with termination factors reveals the translocated cricket paralysis virus IRES. Mol Cell 57(3):422–432.

20 Kerr CH, Ma ZW, Jang CJ, Thompson SR, & Jan E (2016) Molecular analysis of the factorless internal ribosome entry site in Cricket Paralysis virus infection. Scientific reports 6:37319.

21 Nakashima N & Uchiumi T (2009) Functional analysis of structural motifs in dicistroviruses. Virus Res 139(2):137–147.

22 Jan E (2006) Divergent IRES elements in invertebrates. Virus Res 119 (1):16–28.

23 Jang CJ, Lo MC, & Jan E (2009) Conserved element of the dicistrovirus IGR IRES that mimics an E-site tRNA/ribosome interaction mediates multiple functions. J Mol Biol 387(1):42–58.

24 Jang CJ & Jan E (2010) Modular domains of the Dicistroviridae intergenic internal ribosome entry site. RNA 16(6): 1182–1195.

25 Hatakeyama Y, Shibuya N, Nishiyama T, & Nakashima N (2004) Structural variant of the intergenic internal ribosome entry site elements in dicistroviruses and computational search for their counterparts. RNA 10(5):779–786.

26 Cevallos RC & Sarnow P (2005) Factor-independent assembly of elongation-competent ribosomes by an internal ribosome entry site located in an RNA virus that infects penaeid shrimp. J Virol 79(2):677–683.

27 Hertz MI & Thompson SR (2011) In vivo functional analysis of the Dicistroviridae intergenic region internal ribosome entry sites. Nucleic Acids Res 39(16):7276–7288.

28 Abeyrathne PD, Koh CS, Grant T, Grigorieff N, & Korostelev AA (2016) Ensemble cryo-EM uncovers inchworm-like translocation of a viral IRES through the ribosome. eLife 5.

29 Zhang H, Ng MY, Chen Y, & Cooperman BS (2016) Kinetics of initiating polypeptide elongation in an IRES-dependent system. eLife 5.

30 Firth AE, Wang QS, Jan E, & Atkins JF (2009) Bioinformatic evidence for a stem-loop structure 5’-adjacent to the IGR-IRES and for an overlapping gene in the bee paralysis dicistroviruses. Virol J 6:193.

31 Ren Q, Au HH, Wang QS, Lee S, & Jan E (2014) Structural determinants of an internal ribosome entry site that direct translational reading frame selection. Nucleic Acids Res 42(14):9366–9382.

32 Wang QS, Au HH, & Jan E (2013) Methods for studying IRES-mediated translation of positive-strand RNA viruses. Methods 59(2):167–179.

33 Dinos G, et al. (2004) Dissecting the ribosomal inhibition mechanisms of edeine and pactamycin: the universally conserved residues G693 and C795 regulate P-site RNA binding. Mol Cell 13(1):113–124.

34 Kozak M & Shatkin AJ (1978) Migration of 40 S ribosomal subunits on messenger RNA in the presence of edeine. J Biol Chem 253(18):6568–6577.

35 Ruehle MD, et al. (2015) A dynamic RNA loop in an IRES affects multiple steps of elongation factor-mediated translation initiation. eLife 4.

36 Kerr CH, et al. (2015) The 5’ untranslated region of a novel infectious molecular clone of the dicistrovirus cricket paralysis virus modulates infection. J Virol 89(11):5919–5934.

37 Yamamoto H, et al. (2016) 70S-scanning initiation is a novel and frequent initiation mode of ribosomal translation in bacteria. Proc Natl Acad Sci U S A 113(9):E1180–1189.

38 Nomura M, Gourse R, & Baughman G (1984) Regulation of the synthesis of ribosomes and ribosomal components. Annu Rev Biochem 53:75–117.

39 Adhin MR & van Duin J (1990) Scanning model for translational reinitiation in eubacteria. J Mol Biol 213(4):811–818.

40 Colussi TM, et al. (2015) Initiation of translation in bacteria by a structured eukaryotic IRES RNA. Nature 519(7541):110–113.

41 Sarabhai A & Brenner S (1967) A mutant which reinitiates the polypeptide chain after chain termination. J Mol Biol 27(1):145–162.

42 Skabkin MA, Skabkina OV, Hellen CU, & Pestova TV (2013) Reinitiation and other unconventional posttermination events during eukaryotic translation. Mol Cell 51(2):249–264.

43 Abaeva IS, Pestova TV, & Hellen CU (2016) Attachment of ribosomal complexes and retrograde scanning during initiation on the Halastavi arva virus IRES. Nucleic Acids Res 44(5):2362–2377.

44 Wills NM, et al. (2008) Translational bypassing without peptidyl-tRNA anticodon scanning of coding gap mRNA. EMBO J27(19):2533–2544.

45 Chen J, et al. (2015) Coupling of mRNA Structure Rearrangement to Ribosome Movement during Bypassing of Non-coding Regions. Cell 163(5):1267–1280.

46 Samatova E, Konevega AL, Wills NM, Atkins JF, & Rodnina MV (2014) High-efficiency translational bypassing of non-coding nucleotides specified by mRNA structure and nascent peptide. Nature communications 5:4459.

47 Pisareva VP, Pisarev AV, & Fernandez IS (2018) Dual tRNA mimicry in the Cricket Paralysis Virus IRES uncovers an unexpected similarity with the Hepatitis C Virus IRES. eLife 7.

48 Paek KY, et al. (2015) Translation initiation mediated by RNA looping. Proc Natl Acad Sci U S A 112(4):1041–1046.

49 Reinganum C (1975) The isolation of cricket paralysis virus from the emperor gum moth, Antheraea eucalypti Scott, and its infectivity towards a range of insect species. Intervirology 5(1-2):97–102.

50 Chtarbanova S, et al. (2014) Drosophila C virus systemic infection leads to intestinal obstruction. J Virol 88(24):14057–14069.

51 Bonning BC & Miller WA (2010) Dicistroviruses. Annu Rev Entomol 55:129–150.

52 Lang BF, et al. (2014) Massive programmed translational jumping in mitochondria. Proc Natl Acad Sci U S A 111(16):5926–5931.

53 Herr AJ, Gesteland RF, & Atkins JF (2000) One protein from two open reading frames: mechanism of a 50 nt translational bypass. EMBO J 19(11):2671–2680.

54 Garrey JL, Lee YY, Au HH, Bushell M, & Jan E (2010) Host and viral translational mechanisms during cricket paralysis virus infection. J Virol 84(2):1124–1138.

55 Krishna NK, Marshall D, & Schneemann A (2003) Analysis of RNA packaging in wildtype and mosaic protein capsids of flock house virus using recombinant baculovirus vectors. Virology 305(1):10–24.

56 Cox J & Mann M (2008) MaxQuant enables high peptide identification rates, individualized p.p.b.-range mass accuracies and proteome-wide protein quantification. Nature biotechnology 26(12):1367–1372.

